# Earl Grey: a fully automated user-friendly transposable element annotation and analysis pipeline

**DOI:** 10.1101/2022.06.30.498289

**Authors:** Tobias Baril, James Galbraith, Alex Hayward

## Abstract

Transposable elements (TEs) are major components of eukaryotic genomes and are implicated in a range of evolutionary processes. Yet, TE annotation and characterisation remains challenging, particularly for non-specialists, since existing pipelines are typically complicated to install, run, and extract data from. Current methods of automated TE annotation are also subject to issues that reduce overall quality, particularly: (i) fragmented and overlapping TE annotations, leading to erroneous estimates of TE count and coverage; (ii) repeat models represented by short sections of total TE length, with poor capture of 5’ and 3’ ends. To address these issues, we present Earl Grey, a fully automated TE annotation pipeline designed for user-friendly curation and annotation of TEs in eukaryotic genome assemblies. Using nine simulated genomes and an annotation of *Drosophila melanogaster*, we show that Earl Grey outperforms current widely-used TE annotation methodologies in ameliorating the issues mentioned above, whilst scoring highly in benchmarking for TE annotation and classification, and being robust across genomic contexts. Earl Grey provides a comprehensive and fully automated TE annotation toolkit that provides researchers with paper-ready summary figures and outputs in standard formats compatible with other bioinformatics tools. Earl Grey has a modular format, with great scope for the inclusion of additional modules focussed on further quality control and tailored analyses in future releases.

## BACKGROUND

Over recent decades, great advances in genome sequencing technologies have accompanied significant decreases in sequencing costs. This has led to a huge increase in the availability of genome assemblies for lineages across the eukaryotic tree of life. Further, the advent of long-read sequencing technologies has resulted in a wealth of high quality and highly contiguous genome assemblies, with chromosome-level resources becoming increasingly common for both model and non-model organisms. The massive availability of high-quality genomic resources for diverse organisms provides an exciting opportunity to further our understanding of eukaryote evolution, and in turn the evolution and diversity of the Transposable Elements (TEs) hosted within eukaryotic genomes. TEs are DNA sequences that are capable of moving from one location to another in a host genome, with their own distinct evolutionary trajectories (McClintock 1956). It is clear that TEs are major components of genome content across the eukaryotic tree of life, and that TEs are a major reservoir of genetic variation (Wells and Feschotte 2020). As such, TEs are implicated in a range of evolutionary processes involved in the generation of genomic novelty, via processes including alteration of coding sequence, chromosomal rearrangements and the modification of gene regulatory networks (Chung et al. 2007; Hof et al. 2016; Chuong et al. 2017; Cosby et al. 2019). The number of host processes in which TEs are implicated continues to grow at a high pace, and the extent to which TEs play a fundamental role in eukaryotic evolution remains an open question.

The complex repetitive nature of TEs previously made their assembly from short reads challenging, as repetitive regions were often collapsed or unresolved in genome assemblies. However, the increasing availability of long-read genomic resources provides a timely opportunity to significantly improve our understanding of TE evolution and diversity through studies that sample genomes at varying scales, from population-wide to kingdom-wide and beyond. However, to do this efficiently and cope with the flood of sequencing data currently being generated, high-throughput automated methodologies are essential.

Tools for annotating the location and identity of TEs within a genome are a basic aspect of genomics, even where there is little ultimate research focus on TE biology. A key example is the assembly of a new genome, where TE annotation is performed so that TEs do not interfere with gene prediction pipelines (Campbell et al. 2014; Kollmar 2019). This involves screening for TEs present in the genome and masking them, so they are filtered out during subsequent steps, typically using annotations that replace their nucleotide sequences with either lowercase letters (softmasking), or another symbol such as ‘N’ or ‘X’ (hardmasking). Given that TE annotation is typically carried out prior to gene prediction, the quality of TE annotation performed can influence the resultant quality of host gene annotation. For example, many TEs contain protein-coding regions that could be incorrectly annotated as host genes (Bourque et al. 2018), while repetitive host gene families can be mistaken for TEs.

TEs are inherently repetitive in nature, and many TE annotation approaches firstly seek to identify ‘families’ of elements composed of closely related TE copies considered to originate from a single initial integration in the genome of interest. TE families are typically defined by thresholds, such that two TE sequences are considered to belong to the same family if they align with at least 80% identity, across at least 80bp and at least 80% of their length (Wicker et al. 2007). Resultant TE families are commonly represented by a single consensus sequence, constructed by aligning multiple copies sampled from the genome that fall within the defined sequence threshold. Collectively, the set of resultant TE models for a given species is referred to as a ‘TE library’. TE families can be identified using ‘library-based’ approaches, which search the genome for similarity to elements already present in TE databases. Alternatively, ‘*de novo*’ analyses use either existing knowledge of TE structure or their repetitive nature to detect TEs within a genome.

One of the most well-known and frequently employed automated TE annotation workflows involves *de novo* TE family curation with RepeatModeler (and more recently RepeatModeler2) (Flynn et al. 2020), followed by annotation of the host genome with the newly created *de novo* library using RepeatMasker (Smit et al. 2013). This approach provides an overview of repeat content, but leaves considerable room for improvement in several areas, including the quality and completeness of TE consensus generation, TE classification, and subsequent host genome TE annotation. The gold standard for TE annotation is manual curation, whereby a *de novo* TE prediction program is run on a genome assembly to identify putative TE sequences, before individual TE copies are extracted and a multiple alignment is generated for each TE ‘family’ (Goubert et al. 2022). These alignments are then curated by eye by a human investigator, which involves identifying hallmarks of TE boundaries (such as target site duplications and long-terminal repeats), extending TE alignments as necessary, or trimming alignments to remove poorly aligned regions unlikely to represent true TE sequence. The strength of manual curation lies in the individual care taken to annotate each TE model. However, the inherent difficulties associated with manual curation methods are not easily surmountable. Manual curation requires expert knowledge of the structure and biology of diverse TE types. It also requires significant time investment and is not a feasible option for general application given the rate that new genomic data are being produced. Consequently, manual curation is generally only applicable to studies focusing on single genomes, or small set of genomes. Furthermore, the individual-based nature of manual curation also leads to reproducibility issues, due to variability introduced when trimming sequence alignments by eye, especially when considering differences in experience, pattern-recognition ability, and human error.

To address the limitations of manual TE curation and facilitate large-scale comparative and population-level studies, a diversity of automated TE annotation tools have been developed (Goerner-Potvin and Bourque 2018). However, many of these tools are species-or element-specific and need a level of technological expertise to install and configure, often requiring multiple dependencies and databases, which can be difficult to locate and apply due to dead weblinks, a lack of updates, and retirement of codebases/dependencies. Compounding these problems is a lack of focus on user-friendliness in the usage processes, which frequently includes issues relating to data compatibility with downstream tools, necessitating rounds of data reformatting or cleaning. Overall, the above issues can act as a significant barrier to discourage or prohibit interested biologists from performing research on TEs.

Automated TE annotation remains a computationally challenging process. This is due to the great diversity and complexity of TE sequences, the huge number of TEs that likely remain uncharacterised, and the gradual post-integration erosion of TE sequences due to host mutational processes, which reduces the ability to recognise and identify them.

Consequently, automated TE annotation tools often struggle to accurately identify TE sequences. Methodologies that generate *de novo* TE consensus sequences often produce TE consensus libraries with high levels of redundancy, where a single TE is represented by multiple TE consensus models, and these models often perform poorly in their designation of TE boundaries (Rodriguez and Makałowski 2022). Following consensus generation, TE annotation can also lead to challenges that affect both *de novo* and library-based methods. Fragmented TE annotations are often inferred, where a single TE is represented by multiple separate annotations instead of a single continuous annotation. In addition, TE annotations often contain overlaps, where a single base pair is annotated with multiple TE identities. Such issues can result in distorted TE counts and coverage estimates, with knock-on implications for downstream analyses. Many of these issues are particularly acute when considering non-model organisms that are distantly related to model reference species, where considerable effort has been expended to provide high quality TE curation.

The release of new genome assemblies has outpaced detailed efforts to characterise their associated TEs. Furthermore, this situation will accelerate with the progress of massive-scale genome sequencing efforts such as the Darwin Tree of Life Project and the Earth Biogenome Project (Lewin et al. 2018), which seek to provide high quality chromosomal-level genome assemblies for all eukaryotic life in the British Isles and planet Earth respectively. Such projects offer huge opportunities to expand our understanding of TE biology. However, the scale of genomic resources available also greatly increases the need for rapid, robust, repeatable, and user-friendly automated TE annotation and analysis approaches, so that genomes are accurately annotated for TEs, and public TE databases are not overwhelmed with poor quality sequences uploaded from substandard annotations.

Here, we present Earl Grey, a fully automated TE annotation pipeline combining widely used library-based and *de novo* TE annotation tools, with TE consensus and annotation refinements, aimed at generating high-quality TE libraries, annotations, and analyses for eukaryotic genome assemblies.

## IMPLEMENTATION

To make Earl Grey as user-friendly as possible, we have provided several solutions to suit the needs of most researchers. Earl Grey is available from a Github repository (https://github.com/TobyBaril/EarlGrey), as a package in Bioconda (Grüning et al. 2018) (recommended installation), as a preconfigured Docker/Singularity container (https://biocontainers.pro/tools/earlgrey), and in a web browser via gitpod (https://gitpod.io). It can be run on Linux distributions, such as Ubuntu, and installed on a local system or HPCs supporting Conda, Docker, or Singularity. Earl Grey is parallelised and makes use of multiple CPU threads to reduce runtime.

Earl Grey runs in a configured environment to avoid conflicts between tool versions and to streamline the installation procedure. Users who do not have RepeatMasker (Smit et al. 2013) and RepeatModeler2 (Flynn et al. 2020) installed, and who do not wish to do this, can install Earl Grey using the Bioconda package or via interactive containers with Docker and Singularity, which will install and configure dependencies automatically and provide a virtual machine in which to perform all Earl Grey analyses. For installation directly from GitHub, it is necessary to have RepeatMasker and RepeatModeler2 pre-installed.

Instructions for the installation of dependencies are provided for users who do not currently have them installed. In an effort to maintain Earl Grey as open-source, Earl Grey has been tested, and is recommended to be used, with the curated subset of the Dfam database of repetitive DNA elements (Hubley et al. 2016) (tested with all versions from release 3.6 onwards: earlier versions are not compatible due to the transition of RepeatMasker libraries to the H5 format). If users have access to RepBase RepeatMasker edition libraries (Jurka et al. 2005; Kapitonov and Jurka 2008), they can also choose to configure RepeatMasker with these in addition to Dfam.

Once installed and configured, Earl Grey will run on a given input genome assembly in FASTA format with a single command. Prior to running Earl Grey, the ‘earlgrey’ conda environment must be activated ‘conda activate earlgrey’. Once the conda environment is active, Earl Grey can be called with the command ‘earlGrey’. There are 3 required options, and 6 optional parameters (Table 1). For example, a run on the *Homo sapiens* genome, with the genome located in the current directory, could be started with the minimum parameters: ‘earlGrey-g homoSapiens.fasta-s homoSapiens-o./homoSapiens_outputs/’. If users require the curation of a TE library for a polyploid species, we recommend providing Earl Grey with a haploid genome assembly for TE library construction, followed by annotation of the full polyploid assembly.

**Table 1.** Parameters for Earl Grey.

| Command Flag | Description | Required? |
| --- | --- | --- |
| -g | Path to genome FASTA file | Y |
| -s | Name for results files (cannot contain spaces) | Y |
| -o | Path to output directory (. represents current directory). Relative file paths can be used. | Y |
| -r | RepeatMasker species search term used for the optional initial mask of | N |
|  | known elements, in string ("Arthropoda") or NCBI taxonomy ID (6656) format. |  |
| -l | User-provided custom TE library in FASTA format for the optional initial mask of known elements. | N |
| -t | Number of threads (default: 1) | N |
| -c | Should the *de-novo* consensus sequences be clustered to reduced library redundancy? (yes\|no, Default: no) | N |
| -m | Do you want to remove all potentially spurious TE annotations <100bp? (yes\|no, Default: no) | N |
| -d | Generate a softmasked genome after annotation? (yes\|no, Default: no) | N |
| -i | Number of iterations to run the BLAST, Extract, Align, Trim process (Default: 10) | N |
| -f | Number of flanking bases to add in each round of the BLAST, Extract, Align, Trim process (Default: 1000) | N |
| -n | Maximum number of sequences to use during consensus improvement during | N |
|  | the BLAST, Extract, Align, Trim process<br>(Default: 20) |  |

Earl Grey runs through a multi-step TE curation and annotation pipeline to annotate a given genome assembly with all intermediate results saved in their respective directories (Figure 1). Logs are printed to ‘stdout’ (the console) and saved to a log file in the Earl Grey output directory. The steps involved in the Earl Grey TE annotation procedure are outlined below:

1. The first step of Earl Grey prepares the input genome for analysis. Some tools used in the pipeline are sensitive to long header names. To prevent associated issues, header names are stored in a dictionary and replaced with generic headers using the naming convention ‘ctg_n’, where ‘n’ is a unique integer for each entry in the FASTA file. Ambiguous nucleotide IUPAC codes are replaced with “N” due to incompatibility with some tools, including the search engines used by RepeatMasker. The original input genome file is backed up and compressed, with the prepared input genome version saved under the same file name appended with the extension ‘.prep’.
2. If required by the user, known repeats are identified and masked using RepeatMasker and a user-specified subset of the TE consensus libraries (ie Dfam and/or RepBase depending on RepeatMasker configuration), or a user-supplied custom FASTA file of consensus sequences. This is a non-default option, as in most cases it is expected that annotation is being performed on a previously unscreened species and a new search is required, rather than grouping TE families with previously described elements. A sensitive search is performed ignoring small RNA genes ‘-s-norna’, and a hard masked version of the input genome is produced with nucleotides within known TEs replaced with ‘N’.
3. The input genome, or masked genome if step 2 is performed, is analysed with RepeatModeler2 for *de novo* TE identification (Flynn et al. 2020). The optional LTR identification step included as part of RepeatModeler2 is not used, as we implement a separate LTR curation step later in the Earl Grey pipeline during the RepeatCraft stage, as this is a requirement for RepeatCraft (see step 12). RepeatModeler2 outputs a library of *de novo* consensus sequences using the following naming convention: “rnd-n_family-n#TE_Classification” (e.g rnd-1_family-256#LINE/R2-Hero), where rnd stands for the RepeatModeler analysis round in which an element was identified (the default of 6 rounds are run in Earl Grey. If annotating very large genomes, manually increasing RepeatModeler2 sampling rounds is recommended).
4. The success of the RepeatModeler2 run is verified as failures can occur when annotating certain genome assemblies. For example, when annotating a genome where enough unsampled nucleotides remain to initiate a new round of RepeatModeler2, but where there are not enough unsampled long sequences for the additional round to run successfully, this leads to a program failure (e.g. https://github.com/Dfam-consortium/RepeatModeler/issues/118). If this occurs, Earl Grey will automatically restart the RepeatModeler2 run with a reduced maximum stage number to ensure it runs successfully.
5. To generate maximum-length *de novo* TE consensus sequences, the *de novo* TE library undergoes an automated implementation of the “BLAST, Extract, Extend” (BEE) process described by (Platt et al. 2016). In Earl Grey, this process is termed “BLAST, Extract, Align, Trim” (BEAT), and is adapted from TEstrainer (https://github.com/jamesdgalbraith/TEstrainer). BEAT is iteratively performed on each consensus sequence until either the sequence cannot be considerably improved (based on increases in sequence length), or, to prevent infinite loops, the maximum number of specified iterations is complete. The default parameters used throughout BEAT (including extending flanks by 1000bp, a maximum of 10 iterations, and the use of 20 sequences for consensus construction) were selected through extensive manual testing (Discussed in additional file 1). However, all parameters can be specified by the user with command line options. Using default options, the BEAT process proceeds as follows: (i) Each iteration begins with a search for tandem repeats within the consensus sequence using Tandem Repeat Finder (TRF) (Benson 1999). If TRF identifies >90% of the consensus sequence as tandem repeats, the consensus is classified as a tandem repeat and not curated further. If 50% to 90% of the consensus sequence is identified as tandem repeats, the longest section not identified as tandem repeats is extracted and treated as the final consensus sequence, and the sequence is not curated further. If TRF finds <50% of the consensus sequence to be tandem repeats, curation proceeds. (ii) Genome-wide copies of the repeat are identified using BLAST+ (Camacho et al. 2009) (-task dc-megablast). The top 20 matching sequences (defined as having >70% pairwise identity and >50% coverage of the query consensus sequence), are selected and both flanks of each of the 20 sequences is extended by 1,000bp. If at any point of the iterative process filtering reduces the number of genome-wide matches to below three, the iteration’s starting consensus sequence is considered as high quality as possible, and further iterative curation ceases. To ensure the selected sequences improve on the original RepeatModeler sequence, the extended sequences are aligned to the initial RepeatModeler consensus sequence using BLAST (-task dc-megablast), and any which do not align with >70% pairwise identity and >50% coverage are discarded. Manual testing found that, without this step, it is possible for the final consensus sequence to occasionally represent a more abundant repetitive sequence that neighbours a copy of the initial RepeatModeler consensus sequence. (iii) The extended sequences are pairwise aligned to each other using BLAST (-task dc-megablast). Using these alignments, the flanking regions of the extended sequences are trimmed to only include sequence which aligns to at least one other extended sequence. A multiple sequence alignment (MSA) of the trimmed, extended sequences and the starting consensus sequence is constructed using MAFFT (Katoh and Standley 2013). Any columns of the resultant multiple sequence alignment (MSA) which contain only a single nucleotide are removed from the MSA, followed by removal of the starting consensus sequence, so that only the new extended and trimmed sequences remain. This first MSA is realigned with MAFFT and a new consensus sequence is constructed using majority rule. Manual testing identified fewer ambiguous positions in consensus sequences constructed from realigned MSAs compared to consensus sequences constructed from the first MSA. This new consensus sequence is aligned to the previous iteration’s consensus sequence using BLAST (-task dc-megablast) to verify if the sequence is improved. Any new sequence either shorter than, or having less than 80% coverage of, the previous consensus sequence is treated as a reduction in quality, and the previous iteration’s consensus sequence is treated as the final consensus sequence. New consensus sequences longer than the starting sequence by <50% of the flank extension size are treated as complete and not curated further. New consensus sequences longer than the previous consensus sequence by >50% of the flank extension size are submitted for a further round of curation. (iv) Following the iterative BEAT curation process, TRF (Benson 1999), MREPS (Kolpakov et al. 2003) and SA-SSR (Pickett et al. 2016) are used to identify potential satellite repeats in the BEAT curated consensus sequence library. Any consensus sequences identified as >50% tandem repeats are considered likely to be simple or satellite repeats. Within this set, any sequences identified as being >90% a contiguous tandem repeat with a period of 200bp or more are classified as macrosatellites, and the consensus is trimmed to a single period of the repeat.
6. If clustering is specified by the user, the set of *de novo* consensus sequences are clustered using cd-hit-est (Li and Godzik 2006; Fu et al. 2012) with parameters satisfying the TE family definition of Wicker et al. (2007), implemented as described by Goubert et al. (2022), i.e. ‘-d 0 -aS 0.8 -c 0.8 -G 0 -g 1 -b 500 -r 1’. The motivation behind clustering is to reduce the presence of redundant TE consensus sequences, such as sequences on opposite DNA strands being considered as separate TEs, or multiple consensus sequences being formed from older fragments of a recognisable TE, which are not recognised as the same element in RepeatModeler. However, we suggest that clustering should be used with caution. For example, TEs can transpose into each another, creating new transposition-competent chimeric TEs, which may or may not continue to mobilise as a unit. In such cases, clustering could result in the TE library containing a single consensus for the longest of similar consensus sequences. Consequently, the individual independently mobilising TEs would be annotated as smaller fragments of the large chimeric TE only, and not as separate elements. Conversely, if clustering is not performed, the resultant TE library will contain a consensus sequence for the chimeric TE, as well as sequences for the individual constituent TEs. Additionally, clustering will also mask highly similar TE subfamilies to leave a single representative, which could impact estimates of TE divergence making some TEs appear older than they would if separate subfamilies were retained. Therefore, the default behaviour is to avoid clustering, but this can be activated at the discretion of the user.
7. If an initial RepeatMasker step was performed, the curated *de novo* consensus sequences are combined with the known TE library subset used during the initial RepeatMasker step to produce a combined TE library. Otherwise, the *de novo* library is used.
8. The *de novo* TE library (or optionally, the combined library, produced in step 7) is used to annotate TEs in the input genome using RepeatMasker, with sensitive search settings (which takes longer, but improves TE detection) and ignoring small RNA genes ‘-s-norna’.
9. The input genome is analysed with LTR_Finder (version 1.07) (Xu and Wang 2007), using the LTR_Finder parallel wrapper (Ou and Jiang 2019), to identify full-length LTR elements, which is required for the TE annotation defragmentation performed in step 10.
10. The TE annotations from the final RepeatMasker run are defragmented and combined with the LTR_Finder results using the loose merge ‘-loose’ process in RepeatCraft (Wong and Simakov 2018) (https://github.com/niccw/repeatcraftp), which produces a modified GFF file containing the refined TE annotations. Briefly, RepeatCraft identifies non-overlapping TE annotations from the same TE family in the same orientation (strand) within 150bp of each other. These annotations are labelled into “groups”, which are then merged to produce a new GFF file with defragmented repeat loci. This improves TE divergence estimates, as remnants of a degraded TE are counted as one insertion rather than several highly-degraded fragments, whilst also reducing inflation of TE copy number.
11. Following defragmentation, Earl Grey removes any overlapping TE annotations using a custom R script (filteringOverlappingRepeats.R, found in the scripts subdirectory of EarlGrey), which assigns half of the length of overlapping segments to each of the two implicated TEs. For example, in the case of a 10bp overlap between two elements, 5bp will be assigned to the first TE, and 5bp will be assigned to the second, thereby ensuring each TE is only annotated and attributed to a single TE.
12. Finally, to decrease the incidence of spurious hits unlikely to be true TE sequences, all TE annotations less than 100bp in length can be removed before the final set of annotated TEs are quantified. This is at the discretion of the user and can be activated with the flag ‘-m yes’.
13. Summary figures are generated. Earl Grey automatically produces the following summary figures providing a general overview of TEs in the input genome (Figure 2): (i) A pie chart illustrating the proportion of the genome assembly annotated with the main TE classifications and non-TE sequence; (ii) A repeat landscape plot, which illustrates the genetic distance between each identified TE and its respective consensus sequence (calculated using the ‘calcDivergenceFromAlign.pl’ utility of RepeatMasker), and is broadly indicative of patterns of TE activity (i.e. recently active TE copies are assumed to have low levels of genetic distance to their respective family consensus). Consistent colour keys are used for the pie chart and repeat landscape to facilitate comparison. At the discretion of the user, a softmasked version of the input genome is generated, with annotated TE sequence replaced with lower-case letters in the resulting FASTA file.
14. Upon completion, the main results are saved within the summary files directory, which contains: (i) TE annotation coordinates in both GFF3 and Bed format; (ii) The TE library used for the final annotation; (iii) High-level summary tables quantifying the main TE classifications, and a family-level summary to show the most abundant TE families; (iv) Summary figures.

**Figure 1.**
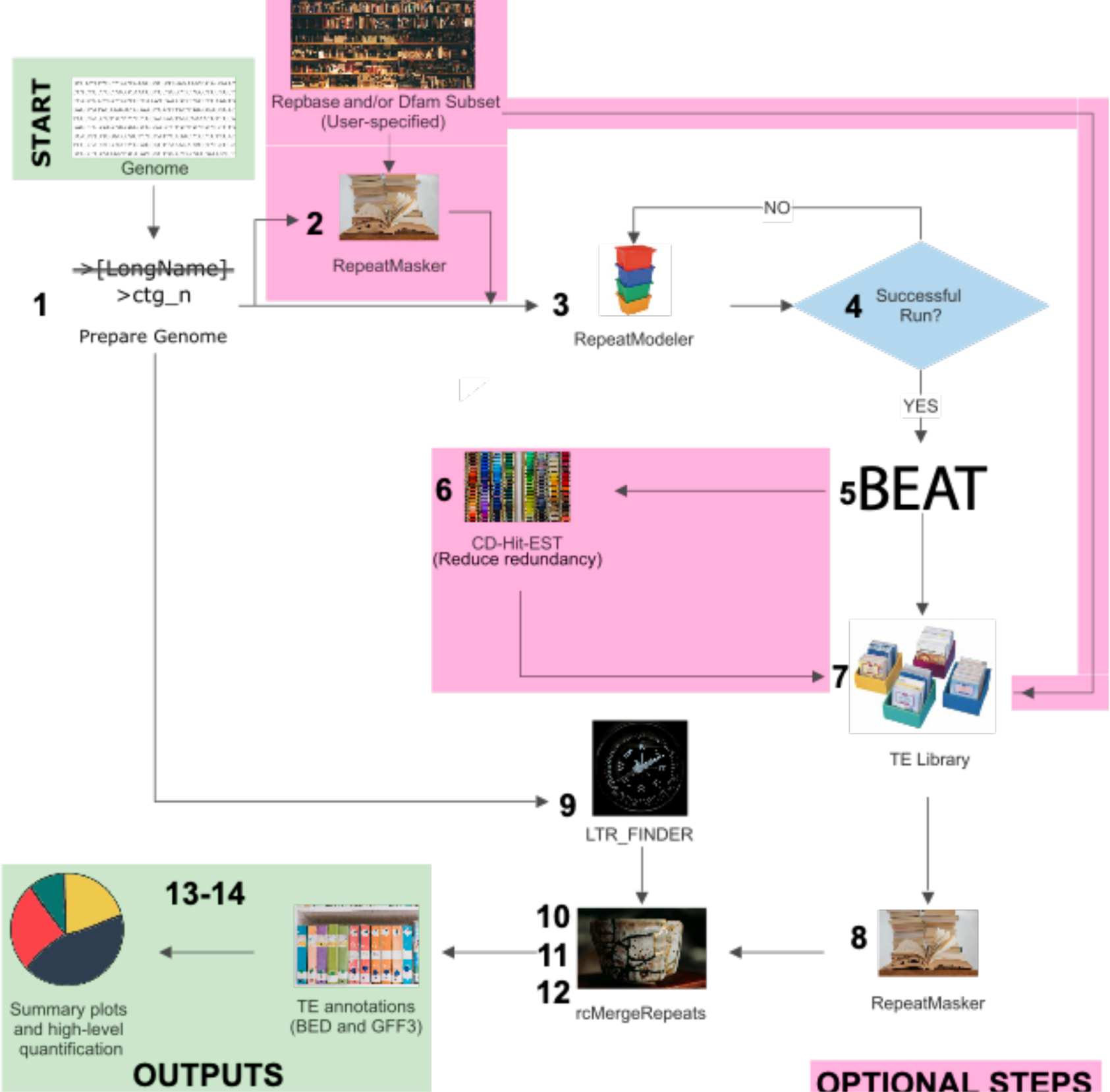
Flow chart illustrating the Earl Grey workflow. Processing is fully automated from start to finish. Stage numbers correspond to steps in the main text.

**Figure 2.**
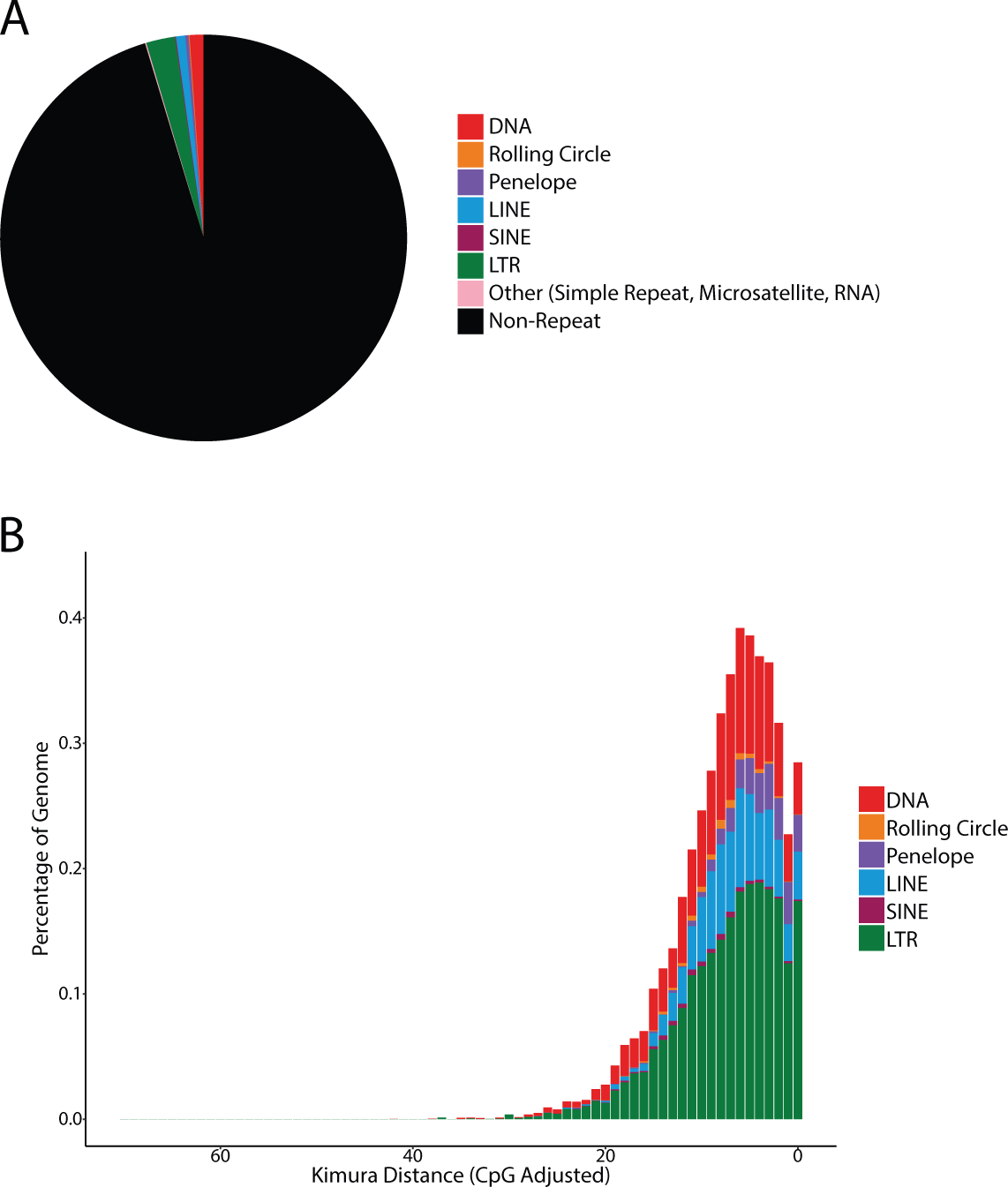
Example summary figures produced by Earl Grey using the results from the annotation of an 840Mb simulated genome with 58% GC content. **A**. Summary pie chart showing the proportion of the genome annotated with the main TE classifications. **B**. Repeat landscape plot, where the X axis indicates divergence from consensus in Kimura distance, and the Y axis indicates the percentage of the genome annotated as TE for each level of divergence. The X axis is reversed relative to the plots produced when using RepeatMasker, such that ancient activity (greater divergence to consensus) appears on the left-hand side, whilst more recent activity is shown towards the right (greater similarity to consensus). Landscape plots can be used to provide an estimate of TE activity within a given genome assembly, with recent activity indicated by a low distance from consensus.

## METHODS

To assess the performance of Earl Grey, we compared it to two widely used existing automated methods: (i) Extensive *de novo* TE Annotator (EDTA) (Ou et al. 2019); (ii) RepeatModeler2 (Flynn et al. 2020) for *de novo* TE consensus generation, followed by RepeatMasker (Smit et al. 2013) for genome assembly annotation. To benchmark Earl Grey against these software, we simulated nine genome assemblies where the coordinates and divergence of all TE copies was known, using scripts from (Rodriguez and Makałowski 2022) (https://github.com/IOB-Muenster/denovoTE-eval). Briefly, this tool is supplied with a configuration file that specifies genomic GC content, TE sequences to be added, TE copy number, expected TE divergence, percentage of TE copies to be fragmented, and percentage of copies to be nested. An initial random DNA sequence of the specified GC content and overall length is simulated. Then, the TE sequence names and copy numbers are taken and random positions in the input genome are assigned to each TE. TE sequences are loaded from a supplied library is FASTA format. These sequences are inserted into the base genome sequence at their assigned positions, taking into account the percentage of copies that should be mutated and fragmented. Subsequently, a second script is run, using the same configuration file, to insert nested TEs into existing TEs added to the base genome in the first round, thereby generating nested insertions. A GFF file is produced containing the coordinates of each TE insertion, the divergence of the copy to the original, whether the copy was fragmented, and whether the copy is nested or not. Configuration files used in this study are supplied in Additional File 2.

We generated the nine simulated genomes in a framework that facilitated consideration of the impacts of both total genome size and extremes of GC content. Genomes were generated with initial sizes (before artificial TE insertion) of 250Mb, 400Mb, and 800Mb, with GC content of 21%, 42% or 58% (Additional File 2). Low and high GC contents were chosen to reflect extremes observed in real genomes. For example, bacterial GC content can vary from <25% to <75% (Hershberg 2016). Fungi also display large variability in GC content, with the rumen fungus *Neocallimastix californiae* having a very low GC content (18.2%) (Peng et al. 2021), whilst the endophytic fungus *Falciphora oryzae* has a high GC content (56.5%) (Xu et al. 2014). Considering sequenced avian, mammalian, and reptilian genomes, GC content tends to lie between 40 and 50% (Bohlin and Pettersson 2019). Therefore, we selected three levels of GC content to reflect TE annotation for the genomes of species from across the tree of life. We inserted 11,883 TE sequences from 30 TE families sourced from Dfam (v3.7) (Hubley et al. 2016; Storer et al. 2021) into each of the simulated genome assemblies, including cut, nested, and diverged copies (up to 30% divergence from consensus). Representatives from a variety of TE classifications were selected, including non-autonomous elements such as MITEs (full configuration datasets for each genome are provided in Additional File 2). TE copy number was determined by generating random numbers following a normal distribution, with the lowest TE copy number being 5 and the highest being 732. This generated a distribution close to what we might expect in a ‘real’ genome assembly, where we anticipate that few TE families would be found at low copy number, the majority at intermediate copy number, and few at very high copy number (Figure 3a). For each simulated genome, the copy number, divergence percentage, nesting percentage, and cutting percentage of each TE family were randomised.

**Figure 3.**
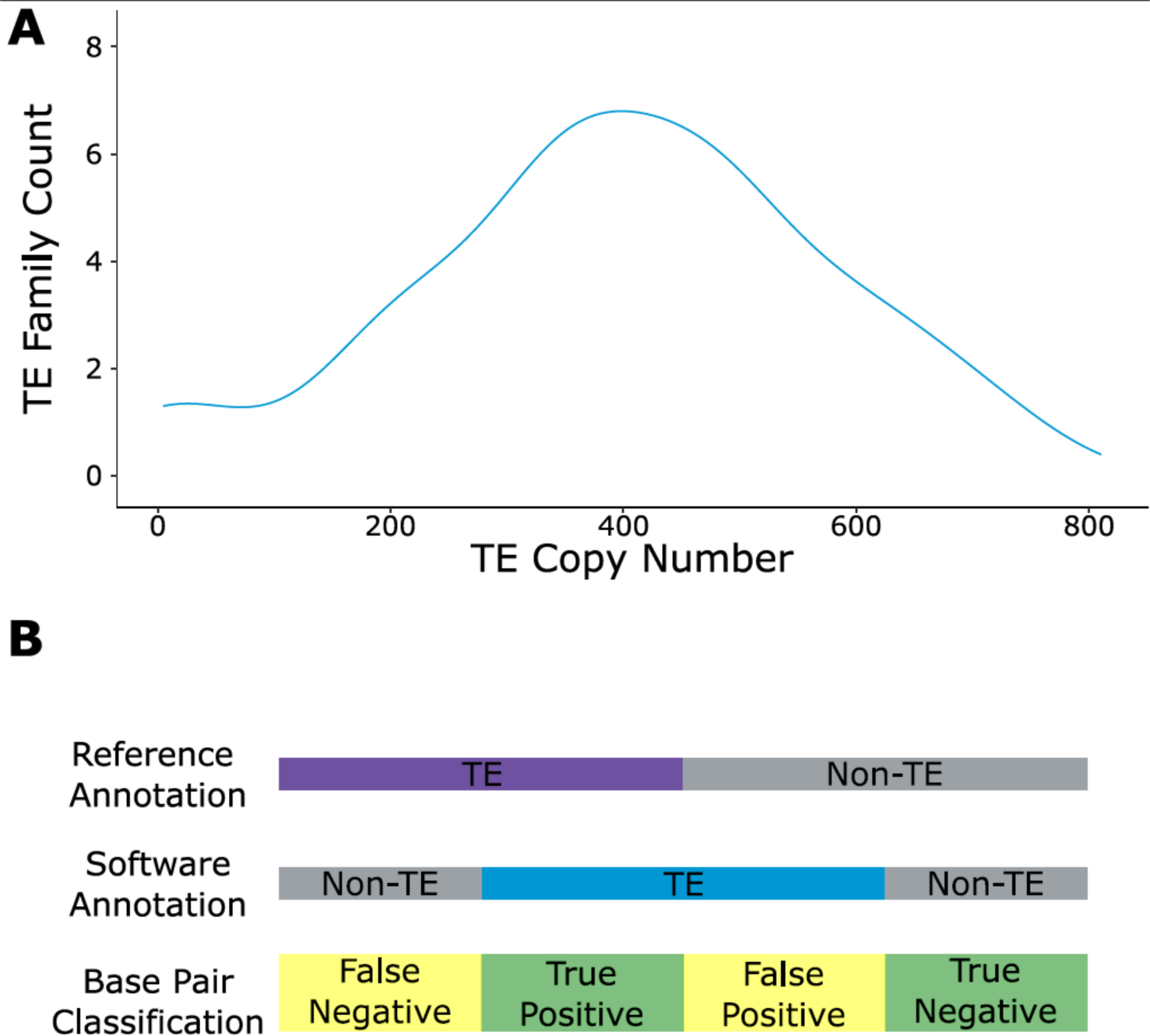
**A**. Distribution showing the number of TE families across copy numbers in the simulated genomes. **B**. Schematic indicating the classification of nucleotides annotated by each method in comparison to their true identity.

The simulated genomes were annotated with EDTA, RepeatModeler2, and Earl Grey using the default options for all tools. For EDTA, the species flag was set to ‘others’, as defined in the documentation for analyses of species other than rice or maize. Using scripts developed and described in (Rodriguez and Makałowski 2022), TE annotation results for each methodology (‘test annotations’) were compared to the ‘reference annotation’ coordinates, detailing the exact position and identity of each TE sequence in the simulated genome, to create a confusion matrix from which the Matthews Correlation Coefficient (MCC) was calculated. An MCC score of +1 arises if all annotations are correct, a score of 0 suggests that test annotations are no better than random guesses, and a score of −1 indicates that all annotations are wrong. Nucleotides were classified based on their agreement between the reference and test annotations as follows: nucleotides found in both reference and test annotations were designated ‘true positive’ (TP), nucleotides absent in both annotations were designated ‘true negative’ (TN), nucleotides found only in the test annotation were designated ‘false positive’ (FP), and nucleotides found only in the reference annotation were designated ‘false negative’ (FN) (Figure 3b) (Rodriguez and Makałowski 2022). TE classifications were compared between the reference and the test annotations using BEDTools intersect and subsequent analyses in R using Rstudio and the tidyverse and ape packages (Paradis et al. 2006; Racine 2013; Team 2013; Wickham et al. 2019). To compare each software-generated TE consensus (query) length with the corresponding real TE consensus (subject) length, the highest-scoring pair for each consensus and real sequence was extracted by filtering for the highest bitscore with BLASTn under default parameters (Camacho et al. 2009). Query and subject total lengths were compared to calculate each consensus’ percentage coverage in comparison to total real TE length.

Following initial benchmarking, the genome assembly of the fruit fly, *Drosophila melanogaster* (Release 6.52) (Strelets et al. 2014) was annotated to test Earl Grey’s performance in a real genomic context.

Overlapping annotations were filtered using a custom R script and GFF files were compared before and after filtering in R using Rstudio and the tidyverse and ape packages (Paradis et al. 2006; Racine 2013; Team 2013; Wickham et al. 2019). Shared and unique TE annotations were identified using BEDTools intersect (-wao) (Quinlan and Hall 2010).

## RESULTS

### Simulated Datasets: Identification of TEs

The performance of Earl Grey was compared to the following widely-used TE annotation methods: (i) Extensive *de novo* TE Annotator (EDTA) (Ou et al. 2019), (ii) RepeatModeler2 (Flynn et al. 2020) for *de novo* TE consensus generation followed by RepeatMasker (Smit et al. 2013) for genome assembly annotation. To assess the relative performance of Earl Grey, we generated nine simulated genomes of varying size and GC content, containing TE insertions extracted from Dfam (see methods). The annotations generated by each pipeline were compared to the real coordinates of each TE insertion in the simulated ‘reference’ genome to create confusion matrices from which the Matthews Correlation Coefficient (MCC) was calculated. A score between +1 and −1 is calculated, where +1 indicates a perfect annotation, −1 indicates a totally wrong annotation, and 0 indicates that the annotation is as good as a random guess. Raw annotation files are provided in additional file 3.

In the simulated genomes, Earl Grey and RepeatModeler2 outperform EDTA, with MCC scores averaging 0.97 and 0.94, respectively (Figure 4). EDTA scored the lowest with an average MCC of 0.67 and has the highest rates of false positive and false negative annotations (Table S1; Additional File 4). Comparing Earl Grey and RepeatModeler2, the difference in average MCC scores comes from Earl Grey’s robust TE annotation in GC-poor (or conversely AT-rich) genomes, where Earl Grey averaged an MCC score of 0.98, whilst RepeatModeler2 averaged 0.84. Considering mid to high GC content, Earl Grey and RepeatModeler2 both score highly (Figure 4, Table S1, Additional File 4).

**Figure 4.**
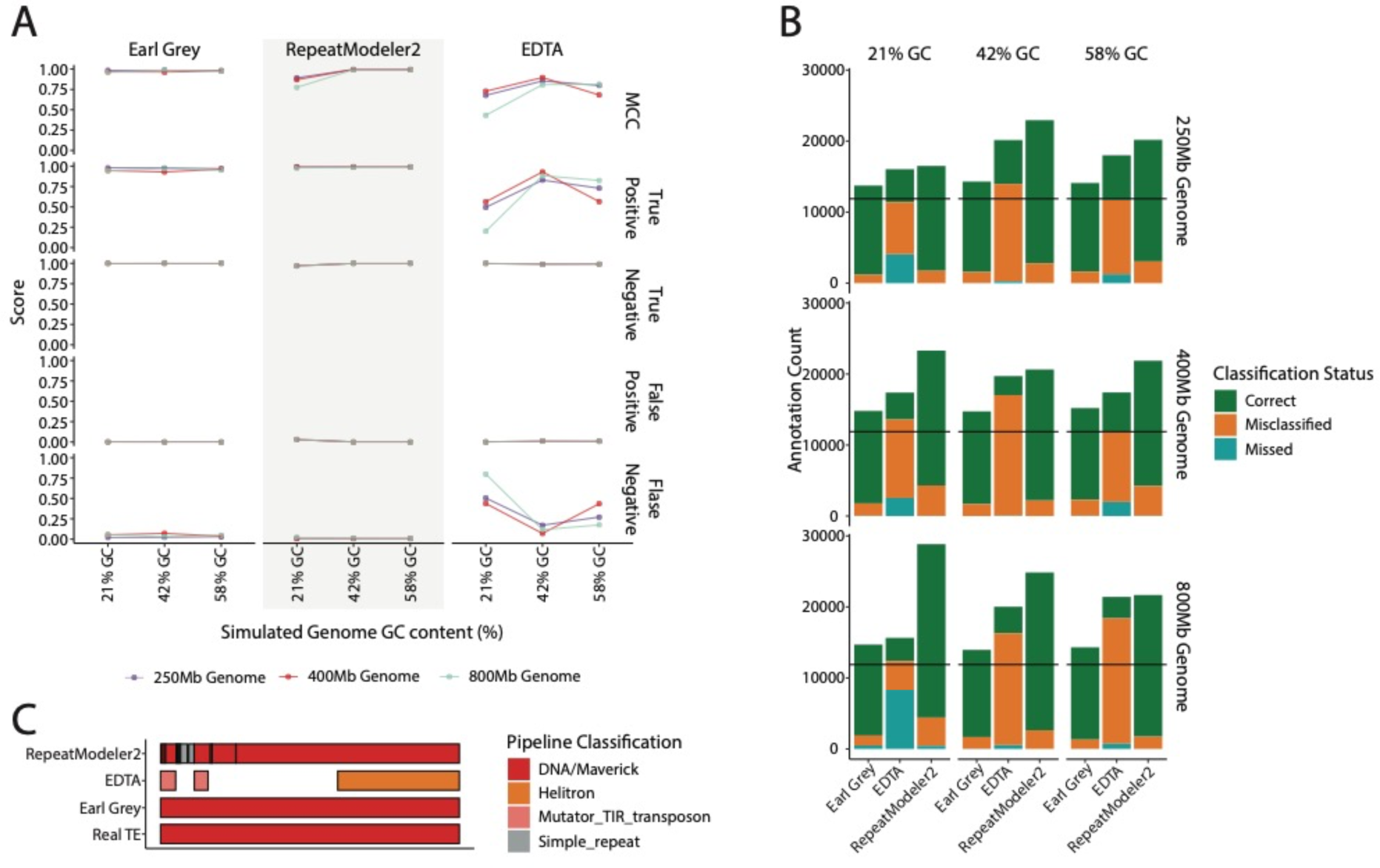
**A**. MCC and nucleotide classification rates for each method compared using the simulated genomes. Colours show genome size as indicated in the key. **B**. Number of TE annotations identified by each method with each annotation labelled as classified correctly, misclassified, or missing when compared to the reference TE annotations. Colours in the key indicate classification status for each TE. The black line indicates the real number of TE insertions. **C.** Schematic demonstrating how the number of TEs annotated can be higher than the real number of TE insertions, as a single TE might be annotated as multiple separate fragments.

Importantly, MCC scores do not consider whether the annotated TE nucleotides were classified correctly. Rather than just identifying that nucleotides are derived from TE sequence as opposed to host genome sequence, it is also important that TEs are correctly classified as the correct TE type. Therefore, we quantified the number of correct, misclassified, and missing TE annotations for each method. Earl Grey had the highest average correct classification (88.5%) whilst RepeatModeler2 averaged 86.6% (Figure 4, Table S2, Additional File 4). EDTA struggled to correctly classify TEs, with a successful classification average of just 23.9% (Figure 4b, Table S2, Additional File 4). Annotations were much more fragmented when using RepeatModeler2, demonstrated by elevated TE counts compared to the number of actual TE insertions, which arises through a single TE being annotated as multiple separate fragments (Figures 4b & 4c).

Overall, Earl Grey performs very well when annotating TEs, with very low false positive and false negative rates and annotations that closely match the real TE loci, leading to very high MCC scores and high correct classification rates. When assessing the small number of TE insertions missed by Earl Grey (an average of 0.47% of real annotations), we find no systematic bias in their classification.

### Simulated Datasets: TE Consensus Libraries

Given Earl Grey’s high performance in correctly annotating TEs, we next examined how Earl Grey’s TE consensus libraries compare to other software that generate *de novo* TE consensus sequences (namely EDTA and RepeatModeler2).

30 real TE families were inserted into the nine simulated genomes. Earl Grey and RepeatModeler2 performed comparably, generating an average of 57.8 and 57.6 consensus sequences, respectively. Earl Grey generated between 49 and 64 consensus sequences, whilst Repeatmodeler2 generated between 47 and 64. EDTA generated the highest number of consensus sequences, with a greatly inflated average of 1,303. EDTA also showed a large range in TE consensus generation, identifying between 21 and 2,948 distinct TE consensus sequences (Table S3, Additional File 4). On average, EDTA identified 796 DNA elements, 169 rolling circle elements, and 338 LTR elements per simulated genome. EDTA did not detect any LINEs, SINEs, or *Penelope-*like elements. Given the high number of consensus sequences generated by EDTA, considering only 30 real TE families were inserted into the simulated genome, we combined the EDTA TE libraries and annotated their consensus sequences with RepeatMasker to interrogate these further. Of the initial 11,727 consensus sequences, 4,903 were annotated with homology to known TEs from Dfam (curated version 3.7) and RepBase (release 20181026). Of these, 3,333 were classified correctly (3,036 LTR, 274 DNA, and 23 rolling circle elements), whilst 1,570 were misclassified, including several that should be classified as non-LTR retroelements (Table S4, Additional File 4). The remaining 6,824 TE consensus sequences generated by EDTA share no similarity to known TEs in Dfam and RepBase, and so we cannot confirm that they represent real TEs. The exclusion of tools to identify non-LTR retroelements is acknowledged in the original EDTA paper: “*Particularly, there is no structure-based program available for the identification of LINEs. The EDTA package may therefore miss a number of elements in, for instance, vertebrate genomes that contain many SINEs and LINEs.*”. The authors suggest the use of RepeatModeler following EDTA annotation. However, this suggestion is not repeated in the current GitHub repository (https://github.com/oushujun/EDTA) or software documentation and may result in researchers missing this suggestion and assuming that EDTA is a complete TE annotation pipeline suitable for analysing diverse genome assemblies.

A key aim of Earl Grey is to generate longer consensus sequences, but without extending them past real TE boundaries. In all simulated genomes, Earl Grey does not generate TE consensus sequences that are significantly different in length from the real reference TE sequences (Table S5, Additional Files 4 and 5). Further, when considering all TE consensus sequences, Earl Grey generates significantly longer consensi than both RepeatModeler2 and EDTA (Table S5, Additional Files 4 and 5), suggesting that these pipelines underestimate TE consensus length, and thus can miss considerable proportions of true TE sequence within a genome. This is further supported when comparing the consensus sequences generated by each method against the reference TE sequences.

Specifically, 88.50% (454/513) of the total consensus sequences generated by Earl Grey are between 95% and 105% of the real TE sequence length, whilst only 21.72% (111/511) of RepeatModeler2 consensus sequences and just 4.49% (23/512) of EDTA consensus sequences are within this length threshold (Figure 5a, Table S6, Additional File 4).

**Figure 5.**
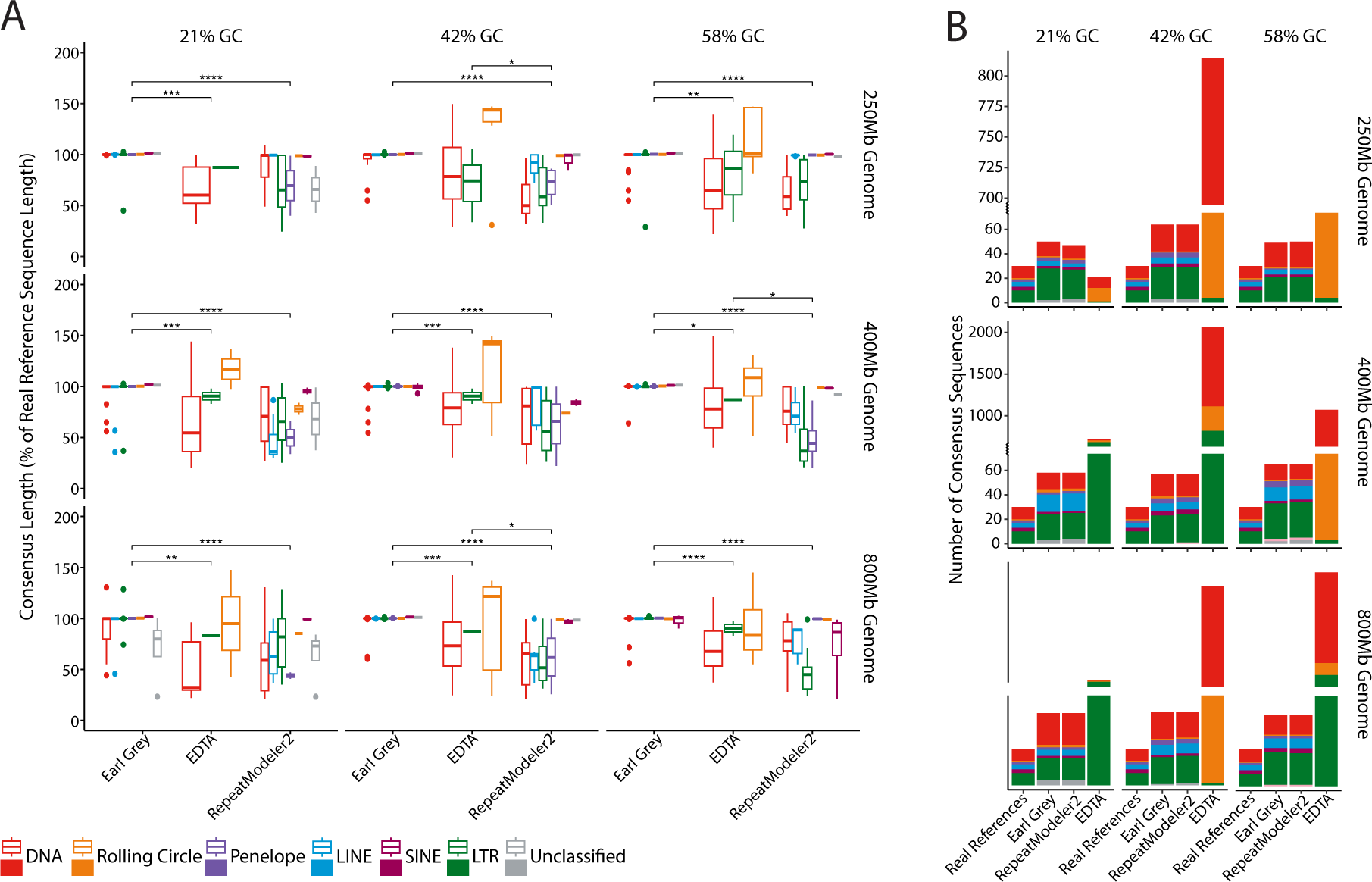
**A**. TE consensus sequence lengths expressed as a percentage of real TE length for each of the main TE classifications. Colours show TE classifications determined by each respective methodology, as indicated in the key. Asterisks indicate levels of significance for comparisons of TE consensus length using Wilcoxon Rank-Sum Tests (*: p≤0.05; **: p≤0.01; ***: p≤0.001; **** p≤0.0001). Only significant comparisons are shown. For readability, Y axis limits were restricted to 200% of real sequence length, removing some data points for EDTA. An extended figure can be found in additional file 6. The DNA TE classification includes DDE-containing TEs, Mavericks/Polintons, and non-autonomous MITEs, as these are labelled as DNA TEs using the RepeatMasker family naming convention (e.g DNA/Maverick and DNA/Tc1-Mariner will both appear in the DNA category). **B**. Number of consensus sequences generated by each methodology, along with the number of real reference sequences. EDTA consensus sequence counts required axis scale adjustment due to the very high consensus sequence number generated. Colours show major TE classifications, as indicated in the key.

Generally, RepeatModeler2 generates TE sequences below the real TE length, with 77.30% (395/511) of RepeatModeler2 consensus sequences below 95% of real TE consensus length. Conversely, EDTA consensus sequences vary considerably in length compared to real consensus sequences, with EDTA generating sequences between 20% and 7,040% of the length of their corresponding real TE sequences, with 68.36% (350/512) being over 105% of the real TE consensus length, and 27.15% (139/512) being <95% of the real TE consensus length (Figure 5a, Table S6, Additional File 4). Earl Grey therefore represents an improvement over RepeatModeler2 and EDTA in approaching full-length TE consensus sequences without significant over-extension past real TE boundaries. Considering individual classifications, Earl Grey generates significantly longer LTR consensus sequences than RepeatModeler2 irrespective of genome size or TE content, whilst generating significantly longer LINE (400Mb 21% and 42% GC; 800Mb 42% GC) and DNA consensus sequences (250Mb 42% GC; 800Mb 42% GC) in certain genomic contexts (Figure 5a, Table S5, Additional File 4). LTR consensus sequences generated by RepeatModeler2 were significantly shorter than the real reference sequences in all simulated genome assemblies except for the 800MB genome with 21% GC content, in which the LINE consensus sequences were significantly shorter (Figure 5a, Table S5, Additional File 4). When compared to EDTA, we find that Earl Grey generates significantly longer LTR consensus sequences (400Mb 21% and 42% GC; 800Mb 21% and 58% GC) (Figure 5a, Table S5, Additional File 4). EDTA LTR consensus sequences were significantly shorter than the real reference sequences in larger simulated genomes (400Mb 21% and 42% GC; 800Mb 21% and 58% GC). EDTA generated significantly longer DNA consensus sequences than RepeatModeler2 in the 800Mb simulated genome with 42% GC content (Figure 5a, Table S5, Additional File 4). However, this result should be taken with caution given the high levels of misclassification observed with EDTA. The generation of longer consensus sequences in Earl Grey can be attributed to the automated implementation of the iterative “BLAST, Extract, Align, Trim” (BEAT) process, that seeks to generate maximum-length consensus sequences from the initial *de novo* TE consensus sequences identified by RepeatModeler2, whilst ensuring consensus sequences do not overextend true TE boundaries.

### Simulated Datasets: TE Annotations

As discussed above, improving the accuracy of TE consensus sequence can have a considerable impact on consensus length, and consequently the amount of correctly annotated TE sequence within a genome. Additionally, Earl Grey was developed to improve several other aspects of TE annotation. Firstly, Earl Grey aims to address current issues with overlapping TE annotations. It is important to address overlapping annotations to prevent inflation of TE count and coverage estimates, as it is not physically possible for a single locus to belong to multiple TEs. When interrogating final annotation outputs, we find overlapping annotations in the outputs of EDTA and RepeatMasker (used to annotate TEs generated with RepeatModeler2), which inflate TE count and coverage estimates. The greatest amount of overlapping annotated TE sequence was found when using EDTA, where removing overlaps reduced overall TE coverage by an average of 7.8% (ranging from 2.4% to 12.7%) (Figure 6a, Table S7, Additional File 4) and TE count by an average of 4.8% (ranging from 1.9% to 8.3%) (Table S7, Additional File 4).

**Figure 6.**
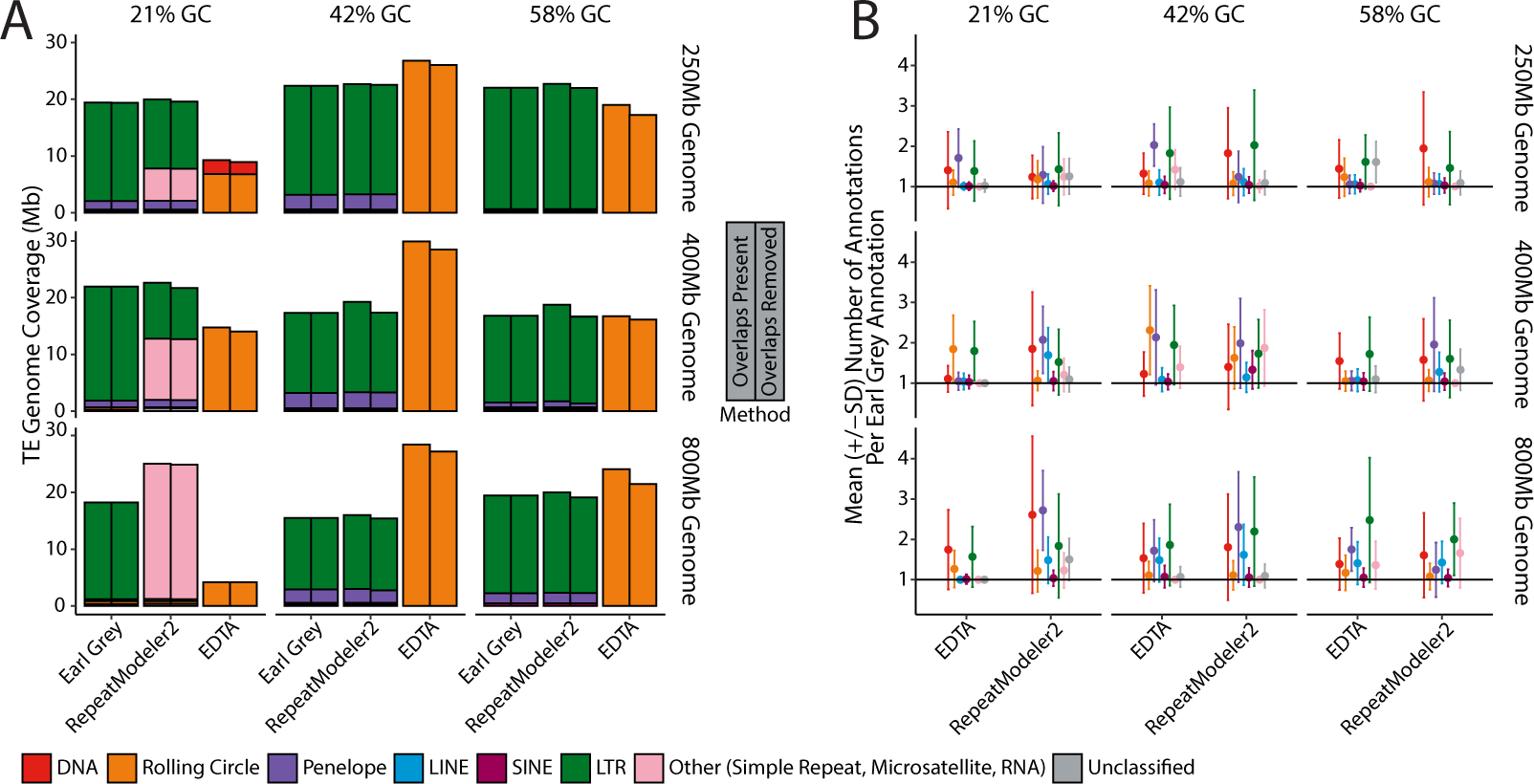
**A**. Number of TEs annotated in the simulated genomes before and after overlapping annotations have been removed. For each method, left hand bars show TE coverage including overlapping annotations, whilst right hand bars indicate TE coverage following removal of overlaps. TE classifications are indicated in the key **B**. Number of annotations per Earl Grey annotation for each simulated genome, demonstrating the level of fragmentation in TE annotation. The black line at 1 indicates the number of annotations expected if annotations are no more fragmented than Earl Grey. TE classifications, as identified by Earl Grey, are indicated in the key.

Secondly, Earl Grey aims to address current issues with the fragmentation of TE annotations, where a single TE can be annotated as multiple fragments, for example due to degradation of the TE sequence. This can inflate estimates of TE copy number and it can also complicate investigations into TE association with host gene features. To address this, Earl Grey includes a post-annotation process employing RepeatCraft (Wong and Simakov 2018) to merge annotations that are likely to belong to the same TE insertion. When examining annotation fragmentation, we find that all methods produce annotations that are more fragmented than Earl Grey, indicated by a mean number of annotations per Earl Grey annotation of >1 for all TE classifications (Figure 6b). Levels of fragmentation were comparable between EDTA and RepeatMasker (using the RepeatModeler2 consensus library), with an average of 1.33 and 1.42 annotations per Earl Grey annotation, respectively (Figure 6b).

### A test using a real genome assembly: TE Consensus Library quality

Following benchmarking of Earl Grey using simulated genomes, we annotated the genome of *Drosophila melanogaster* (Release 6.52) (Strelets et al. 2014) to compare Earl Grey to other software in a real genome assembly. TE annotations from each pipeline are provided in Additional File 7.

Earl Grey and RepeatModeler2 each generated 388 consensus sequences. However, there were fewer unclassified consensus sequences when using Earl Grey (Earl Grey: 88% classified; RepeatModeler2: 81% classified) (Table S8, Additional File 4). Meanwhile, EDTA generated 583 consensus sequences (Table S8, Additional File 4) (Figure 7). With the exception of rolling circle elements, SINEs, and unclassified elements, Earl Grey also generated significantly longer TE consensus sequences (DNA: Kruskal-Wallis, χ^2^_2_ = 21.3, p < 0.01; LTR: Kruskal-Wallis, χ^2^_2_ = 204.0, p < 0.01; LINE: Wilcoxon Rank Sum, W = 5578.5, p < 0.01; Unclassified: Wilcoxon Rank Sum, W = 1919.5, p < 0.01;) (Figure 8). This can be attributed to the automated implementation of the iterative “BLAST, Extract, Align, Trim” process in Earl Grey that works to generate maximum-length consensus sequences from initial *de novo* TE consensus sequences identified by RepeatModeler2. The significantly longer rolling circle consensus sequences generated by EDTA (Kruskal-Wallis, χ^2^_2_ =48.1, p < 0.01) should be treated with caution, due to the very high levels of misclassification observed when using EDTA in the simulated genomes, with these consensus sequences likely instead belonging to a variety of different TE classifications. No significant difference in consensus length between Earl Grey and RepeatModeler2 was found for SINEs and unclassified elements (SINEs: Wilcoxon Rank Sum, W = 2.5, p = 1; Unclassified: Wilcoxon Rank Sum, W = 1919.5, p = 0.15).

**Figure 7.**
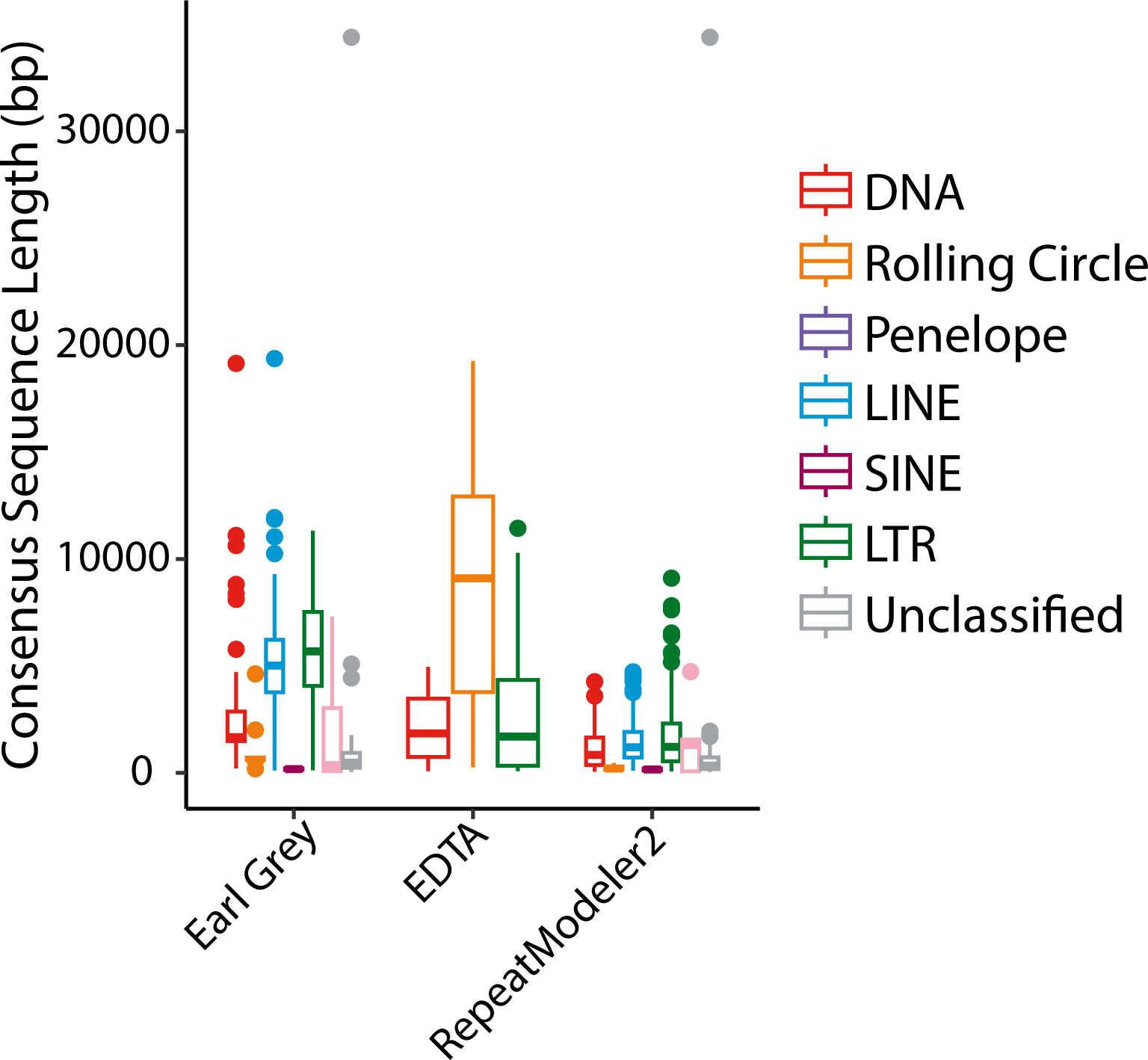
Length of TE consensus sequences split by TE classification and consensus generation methodology. EDTA results should be taken with caution given widespread instances of TE misclassification identified in the simulated genomes. Colours represent TE classifications, as indicated in the key.

**Figure 8.**
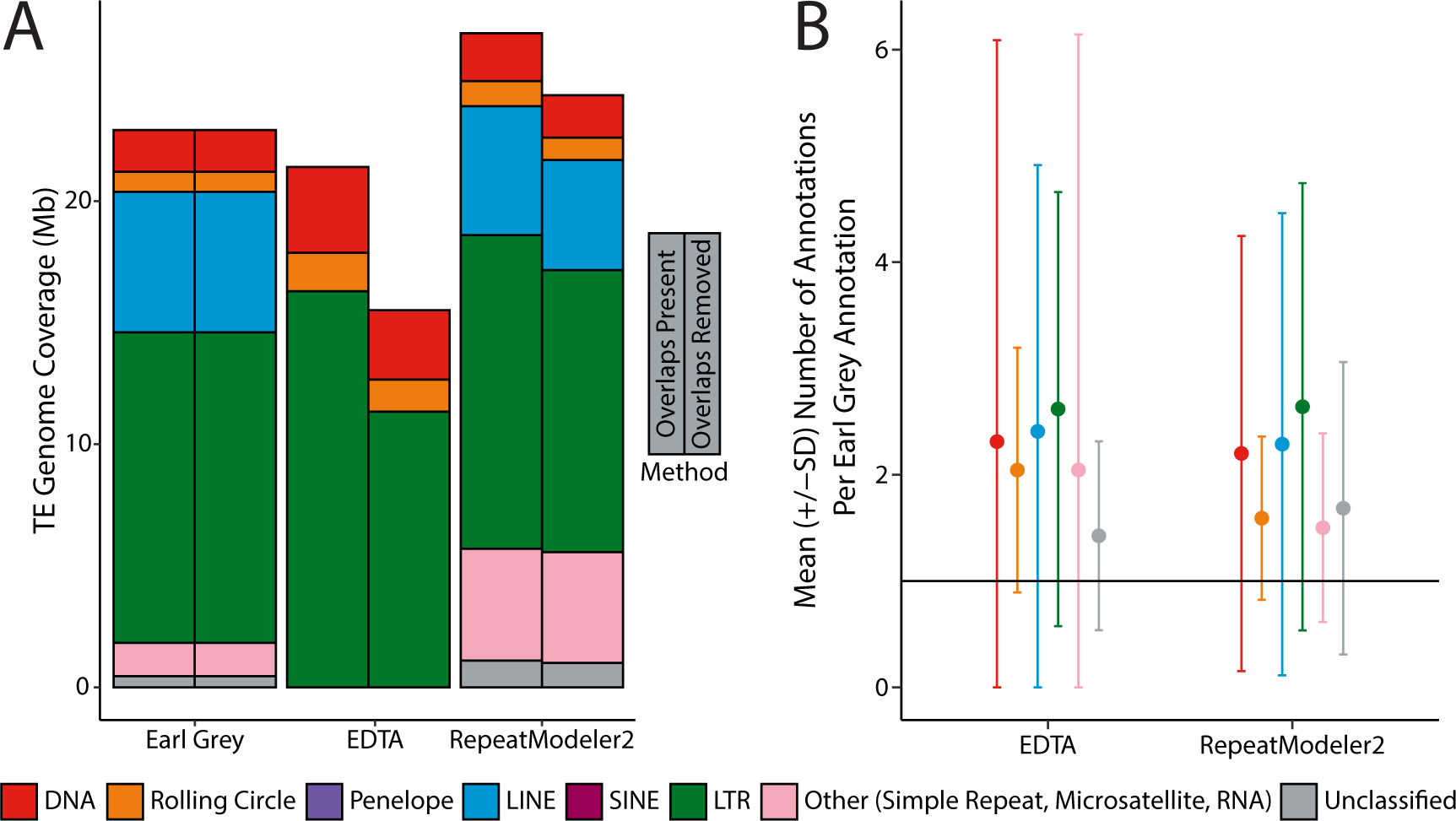
TE classifications are indicated by the colour in the key. **A**. TE coverage expressed as percentage of the total *Drosophila melanogaster* genome assembly annotated using each methodology. Coverage before and after removing overlapping annotations is shown, with TE types coloured as indicated in the key. **B**. Mean number of TE annotations per Earl Grey annotation at shared loci. Error bars indicate 1 standard deviation (SD) from the mean (circles). The black line indicates a perfect match where one annotation matches one annotation using Earl Grey.

### A test using a real genome assembly: TE Annotation quality

When comparing TE annotation results, we find higher TE proportions annotated when using Earl Grey in comparison to EDTA (Figure 8, Table S9; Additional File 4). However, RepeatMasker used with the RepeatModeler2 *de novo* TE library led to higher proportions of the genome being annotated as TE than Earl Grey in *D. melanogaster* (Figure 8, Table S9; Additional File 4).

The highest abundance of unclassified elements was annotated when using RepeatModeler2. This is due to RepeatModeler2 annotating putative TE sequences that do not share sufficient similarity with the TEs in Dfam and RepBase to enable their classification using the RepeatClassifier module. This is surprising given the abundance of knowledge regarding the TE content of *D. melanogaster*, which has been a focus of TE research for at least 43 years (Green 1980; Mérel et al. 2020), and for which substantial TE resources exist. Earl Grey also makes use of the RepeatClassifier module, but it does so with longer and more refined consensus sequences, which likely reveals sufficient homology to known elements to enable their classification.

When interrogating final annotation outputs, we find numerous overlapping annotations in the outputs of EDTA and RepeatMasker (using the RepeatModeler2 library), which inflate TE count and coverage estimates (Figure 8, Table S9, Additional File 4). For EDTA, removing these overlaps reduces total annotation coverage by 22%, whilst annotation coverage is reduced by 9.5% for RepeatMasker (using the RepeatModeler2 library) (Figure 8).

Annotations produced with RepeatMasker (using the RepeatModeler2 library) and EDTA are more fragmented than Earl Grey annotations. This is demonstrated by each Earl Grey annotation, of those that are shared, overlapping with multiple annotation loci generated by the other methodologies (Figure 8). Specifically, each Earl Grey Annotation was represented by an average of 2.14 annotations when using EDTA, and 1.98 annotations when using RepeatMasker (with the RepeatModeler2 library). For all TE classifications, the level of fragmentation was reduced when using Earl Grey, demonstrating Earl Grey’s ability to defragment repeat annotations in a real genomic context.

## DISCUSSION

We have introduced Earl Grey, a fully automated transposable element annotation and analysis pipeline for repeat identification in genome assemblies of diverse organisms. Earl Grey provides various benefits over other pipelines employed for TE annotation. Specifically, Earl Grey was designed to increase TE consensus sequence length, resolve spurious overlapping and fragmented annotations, offer user-directed flexibility in TE annotation parameters, provide users with results in standard formats for compatibility with downstream analyses, and improve the user-friendliness of TE annotation. In addition, “paper-ready” summary figures are produced to provide researchers with a high-level overview of the TE landscape and activity profile for a given genome assembly.

Benchmarking of Earl Grey shows favourable improvements in TE annotation compared to other commonly used pipelines, with an MCC very close to a perfect score of +1. In addition to being robust in terms of TE consensus generation and subsequent TE annotation, Earl Grey will benefit researchers requiring TE annotation in an “all-in-one” automated package requiring no extra analysis tools or steps. Although not as accurate as Earl Grey, RepeatModeler2 scored favourably in benchmarking, but it requires a separate RepeatMasker run following TE library generation to annotate a genome. In addition, Earl Grey is more robust than existing methods when analysing AT-rich genomes, making it suitable for annotation of genome assemblies representing diverse species from across the eukaryotic tree of life.

Earl Grey provides extra polishing steps to further optimise TE annotation results. Through the implementation of the automated “BLAST, Extract, Align, Trim” process, Earl Grey succeeds in producing longer TE consensus sequences without extending significantly beyond true element boundaries (Figure 7b). In the context of TE annotation, the longer TE consensus sequences generated by Earl Grey are beneficial in comparison to the shorter ones produced by the other software, as finding longer matches between individual putative TE copies in a genome brings us closer to confident identification of TE boundaries and more accurate estimations of genome-wide TE content, which remains a significant challenge within the TE field, and is of direct utility for various research applications. There is little chance of the TE boundary sequence being annotated if it is not found in the shorter TE consensus sequences generated by other software. Therefore, finding longer matches among individual putative TE copies in a genome, without extending to generate consensus sequences longer than real insertions, presents a step forward in our ability to accurately identify TEs in genome assemblies. It should be noted that we assessed the performance of RepeatModeler2 under default settings. There is an optional LTR module available for RepeatModeler2, which is likely to improve the performance of RepeatModeler2 for LTR consensus generation in terms of LTR sequence length and completeness.

As previously discussed by (Rodriguez and Makałowski 2022), redundancy in the models generated by existing *de novo* TE identification tools is an issue during annotation. We have addressed this in Earl Grey through the inclusion of optional library redundancy reduction steps. By clustering after the BEAT process, only TE sequences from different putative TE families are included in the final TE consensus library. However, this remains an optional step in the Earl Grey pipeline and should be used with caution, including manual evaluation of redundancies.

When considering the extent of overlap in annotations produced by other methods, Earl Grey demonstrates a significant improvement over existing pipelines by eliminating overlapping TE annotations. This reduces the risk of inflated TE counts whilst remaining accurate in estimating TE coverage in genome assemblies. Furthermore, the removal of overlapping annotations ensures that each base pair of the genome assembly is only annotated as a single TE, so that results remain biologically plausible.

## FUTURE DEVELOPMENTS

A key aim of Earl Grey was to produce a user-friendly TE annotation pipeline that can facilitate large-scale comparative studies through a fully automated process. This first release of Earl Grey provides an improved starting point for automated TE curation and annotation, which can be built upon, to meet the developing needs of the genomics research community. To this end, we have identified several areas of development to be considered for future implementation.

Whilst efforts have been made to improve current automated curation methodologies, we acknowledge that for the foreseeable future, the gold standard will remain manual curation. However, this is also possible following Earl Grey analysis, which can be used to accelerate the process, as demonstrated previously for an in-depth annotation of the Monarch butterfly genome (Baril and Hayward 2022).

Features of the biology of certain TEs can make automatic curation particularly challenging, and several challenges remain. For example, many LINE insertions (and PLE insertions) are heavily 5’ truncated, due to premature termination of reverse transcription during first strand synthesis, either as a consequence of polymerase inefficiency or host cleavage of the RNA intermediate (Suzuki et al. 2009; Baldwin et al. 2024). In situations where there is an accumulation of 5’ truncated elements and the retention of only very few full-length sequences, automated curation can result in a 5’-incomplete LINE consensus sequence, given insufficient numbers of columns with identities in the 5’ regions of the multiple sequence alignments used to compute the consensus (Figure 5a). To address this, future improvements could involve the differential treatment of the 5’ and 3’ ends of putative LINE consensus sequences to allow extension of the 5’ end with fewer column identities in the alignment compared to the 3’ end.

There are inherent limitations with all-by-all genome alignment methods for the identification of repetitive elements. In particular, very low copy number TE families evade detection due to the thresholds used to automatically define ‘repetitive sequences’. If resources are available, some low copy number TE families can be detected using a population or pangenomic approach, which can enable all-by-all comparison algorithms to detect these in several copies, as used to generate a comprehensive TE library for the fungal pathogen, *Zymoseptoria tritici* (Baril and Croll 2023).

Currently, Earl Grey assigns 50% of overlapping base pairs to each of the overlapping TE annotations to resolve overlaps. In an optimal scenario, it would be beneficial to determine which TE each overlapping base belongs to. Whilst this seems like a simple issue to solve on the surface, it actually represents a deeply complex problem that is difficult to solve with existing approaches. Overlaps arise when the bases match with similar scores to competing annotations, meaning that existing algorithms cannot determine which base belongs to which TE. To overcome this, novel approaches are needed that can resolve similar-scoring base pairs to the correct TE. This may involve consideration of inter-classification differences in TE dynamics and degradation patterns to make informed predictions on the likelihood of bases belonging to one TE rather than another. However, this will require significant improvements in our understanding of TE-host dynamics, and a movement away from BLAST-based annotation to approach this problem.

We retain the RepeatMasker TE consensus naming convention in Earl Grey, to maximise compatibility with other commonly-used tools. However, we acknowledge the shortfalls of this system, which have become apparent as understanding of TE evolution and diversity has developed. Consequently, users should be aware of the limitations associated with the current naming convention, and we recommend that they should always consult the full TE classification names when considering TE type. For example, the RepeatMasker “DNA” classification includes DDE-containing DNA TEs, non-autonomous DDE DNA TEs, their much smaller derivative non-autonomous MITEs, and the much larger and evolutionarily divergent Maverick/Polinton TEs. Further, *Penelope*-like elements are designated as “LINE/Penelope”, despite their different transposition mechanism.

A major challenge in identifying *de novo* TEs in genome assemblies concerns the annotation of non-TE sequences. Many *de novo* methods, including those employed by Earl Grey, work by identifying sequences found in multiple copies in the genome. As TE annotation is often performed prior to gene annotation, care should be taken that multicopy genes are not incorrectly designated as TE sequences. For example, this can occur when annotating TEs in animals, where olfactory receptor genes are frequently duplicated and exist in expanded copy number families (Mombaerts 1999). To overcome this, gene annotations or models can be used to retain genomic loci known to contain these genes of interest prior to TE annotation, although this can only be done if such models exist, and prior knowledge is held. To address this, we plan to develop a module that will identify multicopy genes and prevent them ending up in TE consensus libraries.

The advancement of genome sequencing technology has been accompanied by a corresponding decrease in sequencing costs. Consequently, the vast majority of genome assemblies being released today are chromosome-level. For example, the Darwin Tree of Life project aims to release ∼70,000 high-quality *de novo* chromosome-level genome assemblies (https://www.darwintreeoflife.org/). With the release of chromosome-level assemblies comes distinct opportunities to further characterise TE landscapes. To add to the current summary plots generated by Earl Grey, it would be informative to automatically generate karyoplots to show the chromosomal distribution of the main TE classifications across the genome assembly (e.g. see examples in (Li et al. 2020; Baril and Hayward 2022)). As well as providing a visual aid, these can help to identify areas of interest for further interrogation. In addition to visual additions, there are opportunities to present the chromosomal distribution of TEs within a quantitative framework. An additional module for Earl could identify statistically significant hotspots and coldspots for TE insertion, where TEs are found in higher, or lower, densities than expected if TEs were evenly distributed across the host genome (Baril and Hayward 2022). To build on this and enable comparisons across higher taxonomic levels, a metric for “evenness of spread” can be included (Baril and Hayward 2022). This can assess the distribution of TEs within a given context and provide an estimate of how even the observed distribution of TE sequences is, in a cross-comparable manner.

Here, we have presented Earl Grey, a new fully-automated TE annotation pipeline for the annotation of genome assemblies. We have shown that Earl Grey improves upon current widely-used TE annotation methodologies in terms of consensus generation, removing overlapping annotations, and reducing fragmentation of annotations. Earl Grey will help to facilitate large-scale comparative genomic studies, as well as being user-friendly and producing outputs in common formats compatible with downstream analyses. There remains scope for further improvements of Earl Grey, especially via incorporating suggestions and feedback from the needs of the research community. Finally, Earl Grey is an open-source project hosted on GitHub. Our aim is for the TE community to request and contribute new features and improvements to the Earl Grey project, so that it is a community-led effort to improve TE annotation.

## AVAILABILITY AND REQUIREMENTS

Project Name: Earl Grey

Project Home Page: https://github.com/TobyBaril/EarlGrey

Operating Systems: Linux-based Systems (e.g Ubuntu), MacOS-based systems (OSX), or a web browser (for the gitpod implementation)

Programming Language: Pipeline including software coded in Python, R, Bash, and Perl Other Requirements: Anaconda3, Docker (Optional), Singularity (Optional)

License: Open Software License v 2.1

Restrictions for Non-Academic Users: None. Some dependencies may require licences for use by non-academic users.

## ABBREVIATIONS

TE: Transposable element

Non-LTR: Non-long terminal repeat LTR: Long terminal repeat

LINEs: Long INterspersed Elements SINEs: Short INterspersed Elements

PLEs: *Penelope*-like Elements

## DECLARATIONS

### Ethics approval and consent to participate

Not Applicable.

### Consent for publication

Not Applicable.

### Availability of Data and Materials

All data generated or analysed during this study are included in this published article and its supplementary information files, which are hosted on Zenodo under the DOI:10.5281/zenodo.10683832 (https://zenodo.org/doi/10.5281/zenodo.10683832)

## Authors Contributions

TB developed Earl Grey, performed analysis, produced the figures, and drafted the manuscript. JG developed the ‘BLAST, Extract, Align, Trim’ scripts for Earl Grey. AH conceived and coordinated the study and participated in writing the manuscript. All authors read and approved the final manuscript.

## Acknowledgements

We thank Dirk-Jan van Workum for assisting with the conda recipe for Earl Grey, users of early iterations of the software for constructive feedback and suggestions, and the reviewers for their constructive comments.

For the purpose of open access, the author has applied a ‘Creative Commons Attribution (CC BY) licence to any Author Accepted Manuscript version arising.

## Additional Files

Additional File 1: Results and discussion for the selection of the default parameters for the BLAST, Extend, Align, Trim (BEAT) process.

Additional File 2: Tar archive containing configuration files and TE families to generate all nine simulated genomes.

Additional File 3: Tar archive containing reference TE coordinates in GFF format and GFF annotation files for each software used to annotate TEs in the simulated genomes.

Additional File 4: Excel file containing all supplementary tables with contents page.

Additional File 5: Distributions of TE consensus lengths generated by each methodology against the real TE consensus sequence lengths.

Additional File 6: Extended version of figure 5a to illustrate the distribution of consensus lengths for EDTA, which extend beyond the limits set for readability.

Additional File 7: Tar archive containing TE annotations for *D. melanogaster* annotated with each software, in GFF format.

## REFERENCES

Baldwin ET, van Eeuwen T, Hoyos D, Zalevsky A, Tchesnokov EP, Sánchez R, Miller BD, Di Stefano LH, Ruiz FX, Hancock M, et al. 2024. Structures, functions and adaptations of the human LINE-1 ORF2 protein. Nature [Internet] 626:194–206. Available from: 10.1038/s41586-023-06947-z

Baril T, Croll D. 2023. A pangenome-guided manually curated library of transposable elements for Zymoseptoria tritici. BMC Res. Notes [Internet] 16. Available from: https://bmcresnotes.biomedcentral.com/articles/10.1186/s13104-023-06613-7

Baril T, Hayward A. 2022. Migrators within migrators: exploring transposable element dynamics in the monarch butterfly, Danaus plexippus. Mob. DNA [Internet] 13:5. Available from: 10.1186/s13100-022-00263-5

Benson G. 1999. Tandem repeats finder: A program to analyze DNA sequences. Nucleic Acids Res. [Internet] 27:573–580. Available from: 10.1093/nar/27.2.573

Bohlin J, Pettersson JH-O. 2019. Evolution of genomic base composition: From single cell microbes to multicellular animals. Comput. Struct. Biotechnol. J. [Internet] 17:362–370. Available from: https://www.ncbi.nlm.nih.gov/pmc/articles/PMC6429543/

Bourque G, Burns KH, Gehring M, Gorbunova V, Seluanov A, Hammell M, Imbeault M, Izsvák Z, Levin HL, Macfarlan TS, et al. 2018. Ten things you should know about transposable elements. Genome Biol. [Internet] 19:199. Available from: 10.1186/s13059-018-1577-z

Camacho C, Coulouris G, Avagyan V, Ma N, Papadopoulos J, Bealer K, Madden TL. 2009. BLAST+: Architecture and applications. BMC Bioinformatics [Internet] 10:1–9. Available from: 10.1186/1471-2105-10-421

Campbell MS, Holt C, Moore B, Yandell M. 2014. Genome Annotation and Curation Using MAKER and MAKER-P. Current Protocols in Bioinformatics [Internet] 48. Available from: 10.1002/0471250953.bi0411s48

Chung H, Bogwitz MR, McCart C, Andrianopoulos A, Ffrench-Constant RH, Batterham P, Daborn PJ. 2007. Cis-regulatory elements in the accord retrotransposon result in tissue-specific expression of the Drosophila melanogaster insecticide resistance gene Cyp6g1. Genetics [Internet] 175:1071–1077. Available from: 10.1534/genetics.106.066597

Chuong EB, Elde NC, Feschotte C. 2017. Regulatory activities of transposable elements: From conflicts to benefits. Nat. Rev. Genet. [Internet] 18:71–86. Available from: 10.1038/nrg.2016.139

Cosby RL, Chang N-C, Feschotte C. 2019. Host–transposon interactions: conflict, cooperation, and cooption. Genes Dev. [Internet] 33:1098–1116. Available from: http://genesdev.cshlp.org/content/33/17-18/1098.full

Flynn JM, Hubley R, Goubert C, Rosen J, Clark AG, Feschotte C, Smit AF. 2020. RepeatModeler2 for automated genomic discovery of transposable element families. Proceedings of the National Academy of Sciences [Internet] 117:9451–9457. Available from: https://www.pnas.org/content/117/17/9451

Fu L, Niu B, Zhu Z, Wu S, Li W. 2012. CD-HIT: accelerated for clustering the next-generation sequencing data. Bioinformatics 28:3150–3152.

Goerner-Potvin P, Bourque G. 2018. Computational tools to unmask transposable elements. Nat. Rev. Genet. [Internet] 19:688–704. Available from: 10.1038/s41576-018-0050-x

Goubert C, Craig RJ, Bilat AF, Peona V, Vogan AA, Protasio AV. 2022. Correction: A beginner’s guide to manual curation of transposable elements. Mob. DNA [Internet] 13:15. Available from: 10.1186/s13100-022-00272-4

Green MM. 1980. Transposable elements in Drosophila and other Diptera. Annu. Rev. Genet. [Internet] 14:109–120. Available from: 10.1146/annurev.ge.14.120180.000545

Grüning B, Dale R, Sjödin A, Chapman BA, Rowe J, Tomkins-Tinch CH, Valieris R, Köster J, Bioconda Team. 2018. Bioconda: sustainable and comprehensive software distribution for the life sciences. Nat. Methods [Internet] 15:475–476. Available from: 10.1038/s41592-018-0046-7

Hershberg R. 2016. Codon Usage and Translational Selection. In: Encyclopedia of Evolutionary Biology. Elsevier. p. 293–298. Available from: 10.1016/b978-0-12-800049-6.00178-5

Hof AEV t., Campagne P, Rigden DJ, Yung CJ, Lingley J, Quail MA, Hall N, Darby AC, Saccheri IJ, van’t Hof AE, et al. 2016. The industrial melanism mutation in British peppered moths is a transposable element. Nature [Internet] 534:102–105. Available from: 10.1038/nature17951

Hubley R, Finn RD, Clements J, Eddy SR, Jones TA, Bao W, Smit AFA, Wheeler TJ. 2016. The Dfam database of repetitive DNA families. Nucleic Acids Res. [Internet] 44:D81– D89. Available from: 10.1093/nar/gkv1272

Jurka J, Kapitonov VV, Pavlicek A, Klonowski P, Kohany O, Walichiewicz J. 2005. Repbase Update, a database of eukaryotic repetitive elements. Cytogenet. Genome Res. [Internet] 110:462–467. Available from: 10.1159/000084979

Kapitonov VV, Jurka J. 2008. A universal classification of eukaryotic transposable elements implemented in Repbase. Nat. Rev. Genet. [Internet] 9:411–412. Available from: 10.1038/nrg2165-c1

Katoh K, Standley DM. 2013. MAFFT multiple sequence alignment software version 7: Improvements in performance and usability. Mol. Biol. Evol. [Internet] 30:772–780. Available from: 10.1093/molbev/mst010

Kollmar M. 2019. Gene Prediction: Methods and Protocols. Humana Press Available from: https://books.google.com/books/about/Gene_Prediction.html?hl=&id=iEkZvwEACAA J

Kolpakov R, Bana G, Kucherov G. 2003. mreps: Efficient and flexible detection of tandem repeats in DNA. Nucleic Acids Res. [Internet] 31:3672–3678. Available from: 10.1093/nar/gkg617

Lewin HA, Robinson GE, Kress WJ, Baker WJ, Coddington J, Crandall KA, Durbin R, Edwards SV, Forest F, Gilbert MTP, et al. 2018. Earth BioGenome Project: Sequencing life for the future of life. Proc. Natl. Acad. Sci. U. S. A. [Internet] 115:4325–4333. Available from: 10.1073/pnas.1720115115

Li W, Godzik A. 2006. Cd-hit: a fast program for clustering and comparing large sets of protein or nucleotide sequences. Bioinformatics 22:1658–1659.

Li Y, Nong W, Baril T, Yip HY, Swale T, Hayward A, Ferrier DEK, Hui JHL. 2020. Reconstruction of ancient homeobox gene linkages inferred from a new high-quality assembly of the Hong Kong oyster (Magallana hongkongensis) genome. BMC Genomics 21:1–17.

McClintock B. 1956. “Controlling Elements and the Gene.” Cold Spring Harb. Symp. Quant. Biol. 21:197–216.

Mérel V, Boulesteix M, Fablet M, Vieira C. 2020. Transposable elements in Drosophila. Mob. DNA [Internet] 11:23. Available from: 10.1186/s13100-020-00213-z

Mombaerts P. 1999. Seven-transmembrane proteins as odorant and chemosensory receptors. Science [Internet] 286:707–711. Available from: 10.1126/science.286.5440.707

Ou S, Jiang N. 2019. LTR_FINDER_parallel: parallelization of LTR_FINDER enabling rapid identification of long terminal repeat retrotransposons. BioRxiv:2–6.

Ou S, Su W, Liao Y, Chougule K, Agda JRA, Hellinga AJ, Lugo CSB, Elliott TA, Ware D, Peterson T, et al. 2019. Benchmarking transposable element annotation methods for creation of a streamlined, comprehensive pipeline. Genome Biol. [Internet] 20:1–45. Available from: 10.1186/s13059-019-1905-y

Paradis E, Strimmer K, Claude J, Jobb G, Opgen-Rhein R, Dutheil J, Noel Y, Bolker B, Lemon J. 2006. ape: Analyses of Phylogenetics and Evolution. *R package version* [Internet] 1. Available from: http://ape-package.ird.fr/ep/diapo_LaReunion_2009.pdf

Peng X, Wilken SE, Lankiewicz TS, Gilmore SP, Brown JL, Henske JK, Swift CL, Salamov A, Barry K, Grigoriev IV, et al. 2021. Genomic and functional analyses of fungal and bacterial consortia that enable lignocellulose breakdown in goat gut microbiomes. Nat. Microbiol. [Internet] 6:499–511. Available from: 10.1038/s41564-020-00861-0

Pickett BD, Karlinsey SM, Penrod CE, Cormier MJ, Ebbert MTW, Shiozawa DK, Whipple CJ, Ridge PG. 2016. SA-SSR: a suffix array-based algorithm for exhaustive and efficient SSR discovery in large genetic sequences: Table 1. Bioinformatics [Internet] 32:2707–2709. Available from: 10.1093/bioinformatics/btw298

Platt RN, Blanco-Berdugo L, Ray DA. 2016. Accurate transposable element annotation is vital when analyzing new genome assemblies. Genome Biol. Evol. [Internet] 8:403–410. Available from: 10.1093/gbe/evw009

Quinlan AR, Hall IM. 2010. BEDTools: A flexible suite of utilities for comparing genomic features. Bioinformatics [Internet] 26:841–842. Available from: 10.1093/bioinformatics/btq033

Racine JS. 2013. RSTUDIO: A PLATFORM-INDEPENDENT IDE FOR R AND SWEAVE. J. Appl. Econometrics [Internet] 27:167–172. Available from: 10.1002/jae

Rodriguez M, Makałowski W. 2022. Software evaluation for de novo detection of transposons. Mob. DNA [Internet] 13:14. Available from: 10.1186/s13100-022-00266-2

Smit AFA, Hubley RR, Green PR. 2013. RepeatMasker Open-4.0. http://repeatmasker.org.

Storer J, Hubley R, Rosen J, Wheeler TJ, Smit AF. 2021. The Dfam community resource of transposable element families, sequence models, and genome annotations. Mob. DNA [Internet] 12:2. Available from: 10.1186/s13100-020-00230-y

Strelets VB, Schroeder AJ, Thurmond J, Goodman JL, Gelbart WM, dos Santos G, Emmert DB, Crosby MA, the FlyBase Consortium. 2014. FlyBase: introduction of the Drosophila melanogaster Release 6 reference genome assembly and large-scale migration of genome annotations. Nucleic Acids Res. [Internet] 43:D690–D697. Available from: 10.1093/nar/gku1099

Suzuki J, Yamaguchi K, Kajikawa M, Ichiyanagi K, Adachi N, Koyama H, Takeda S, Okada N. 2009. Genetic evidence that the non-homologous end-joining repair pathway is involved in LINE retrotransposition. PLoS Genet. [Internet] 5:e1000461. Available from: 10.1371/journal.pgen.1000461

Team RC. 2013. R: A language and environment for statistical computing.

Wells JN, Feschotte C. 2020. A Field Guide to Eukaryotic Transposable Elements. Annu. Rev. Genet. [Internet] 54:539–561. Available from: 10.1146/annurev-genet-040620-022145

Wicker T, Sabot F, Hua-Van A, Bennetzen JL, Capy P, Chalhoub B, Flavell A, Leroy P, Morgante M, Panaud O, et al. 2007. A unified classification system for eukaryotic transposable elements. Nat. Rev. Genet. [Internet] 8:973–982. Available from: 10.1038/nrg2165

Wickham H, Averick M, Bryan J, Chang W, McGowan LD, François R, Grolemund G, Hayes A, Henry L, Hester J. 2019. Welcome to the Tidyverse. Journal of Open Source Software 4:1686.

Wong WY, Simakov O. 2018. RepeatCraft: a meta-pipeline for repetitive element de-fragmentation and annotation. Bioinformatics [Internet] 35:1051–1052. Available from: 10.1093/bioinformatics/bty745

Xu X-H, Su Z-Z, Wang C, Kubicek CP, Feng X-X, Mao L-J, Wang J-Y, Chen C, Lin F-C, Zhang C-L. 2014. The rice endophyte Harpophora oryzae genome reveals evolution from a pathogen to a mutualistic endophyte. Sci. Rep. [Internet] 4:5783. Available from: 10.1038/srep05783

Xu Z, Wang H. 2007. LTR_FINDER: an efficient tool for the prediction of full-length LTR retrotransposons. Nucleic Acids Res. [Internet] 35:W265–W268. Available from: https://www.ncbi.nlm.nih.gov/pubmed/17485477

